# Intrinsic skeletal muscle function and contraction-stimulated glucose uptake do not vary by time-of-day in mice

**DOI:** 10.1101/2024.05.15.594323

**Authors:** Liam S. Fitzgerald, Shannon N. Bremner, Samuel R. Ward, Yoshitake Cho, Simon Schenk

## Abstract

A growing body of data suggests that skeletal muscle contractile function and glucose metabolism vary by time-of-day, with chronobiological effects on intrinsic skeletal muscle properties being proposed as the underlying mediator. However, no studies have directly investigated intrinsic contractile function or glucose metabolism in skeletal muscle over a 24 h circadian cycle. To address this, we assessed intrinsic contractile function and endurance, as well as contraction-stimulated glucose uptake, in isolated extensor digitorum longus and soleus from female mice at four times-of-day (Zeitgeber Times 1, 7, 13, 19). Significantly, while both muscles demonstrated circadian-related changes in gene expression, intrinsic contractile function, endurance, and contraction-stimulated glucose uptake were not different between the four time points. Overall, these results demonstrate that time-of-day variation in exercise performance and the glycemia-reducing benefits of exercise are not due to chronobiological effects on intrinsic muscle function or contraction-stimulated glucose uptake.

**Impact statement:** *Ex vivo* testing demonstrates that there is no time-of-day variation in the intrinsic contractile properties of skeletal muscle (including no effect on force production or endurance) or contraction-stimulated glucose uptake.

## Introduction

Through circadian cycles, or rhythms, time-of-day impacts many aspects of mammalian physiology (Krauchi and Wirz-Justice, 1994; Logan and McClung, 2018; Millar-Craig et al., 1978). Recently, a large focus of the field of exercise physiology has been on the effect of time-of-day on exercise capacity and athletic performance. On the whole, measures of both strength and endurance performance tend to be lower in early morning and higher in the afternoon/evening (Atkinson and Reilly, 1996; Atkinson and Speirs, 1998; Baxter and Reilly, 1983; Bessot et al., 2006; Czelusniak et al., 2021; Douglas et al., 2021; Edwards et al., 2005; Martin et al., 1999; Reilly et al., 2007; Reilly and Down, 1986; Rodahl et al., 1976). While a number of factors have been proposed to underlie these performance impacting effects of time-of-day, including body temperature (Bergh and Ekblom, 1979; Harrison and Bers, 1989), motor unit recruitment (Gueldich et al., 2017; Nicolas et al., 2007; Sedliak et al., 2008), and meal timing/muscle glycogen status (Kerksick et al., 2017; Koch et al., 2020), an emphasized point in the field is that variation in exercise performance is due to circadian fluctuations in the intrinsic properties of skeletal muscle (Douglas et al., 2021). In contrast to this, recent work found that the maximal intrinsic force generating capacity of the mouse extensor digitorum longus (EDL) is not different between two different times of the light phase (i.e. zeitgeber time [ZT] ZT1 and ZT9) (Kahn et al., 2024; personal communication with Dayanidhi S, Kahn RE, Lieber RL). While this study is interesting, whether there are intrinsic changes in skeletal muscle physiology over the course of a 24 h circadian cycle, or in other muscles, remains unknown. Moreover, whether other intrinsic muscle properties, such as submaximal contractile function or resistance to fatigue are impacted across a 24 h circadian cycle, is unknown.

Exercise is a cornerstone therapeutic for preventing or treating clinical hyperglycemia (“Reduction in the Incidence of Type 2 Diabetes with Lifestyle Intervention or Metformin,” 2002;Bajpeyi et al., 2009; Balducci et al., 2006; Cohen et al., 2008; Evans et al., 2005; Loimaala et al., 2003). A fundamental reason for this beneficial effect of exercise is that muscle contraction potently stimulates glucose disposal from the blood into the exercising skeletal muscle (Constable et al., 1988), and does so in an insulin-independent manner (Goodyear et al., 1990; Goodyear and Kahn, 1998; Ploug et al., 1984). Interestingly, like exercise performance, a large focus of the field of exercise physiology, and more broadly the field of diabetes care, has been on the effect of time-of-day on the glucose-controlling benefits of exercise (Riddell et al., 2023). Indeed, in humans, undertaking an exercise training intervention in the afternoon has been found to be superior at improving glycemic control and skeletal muscle insulin sensitivity to exercise training in the morning (Mancilla et al., 2021; Qian et al., 2023; Savikj et al., 2019). Similarly, in mice, a single exercise bout in the early part of the active (i.e., dark) phase, as compared to the early rest (i.e., light) phase, promoted a glycolytic transcriptional signature in skeletal muscle and reduced blood glucose concentration (Sato et al., 2019). Nevertheless, whether this time-of-day effect exercise on glycemic control is due to chronobiological variation in the intrinsic capacity of contraction-stimulated glucose uptake by skeletal muscle has not been studied.

Addressing these major gaps in knowledge, we used an *ex vivo* approach to assess intrinsic contractile function (sub-maximal and maximal) and fatigability, and contraction-stimulated glucose uptake, at four different times of the 24 h cycle (ZT1, ZT7, ZT13, ZT19). An *ex vivo* approach allowed us to study skeletal muscle in isolation, and thus, independent of other factors that might influence contractile function or glucose metabolism (e.g., muscle temperature, blood flow, nerve function, humoral factors, etc.). We also studied two different muscles, the extensor digitorum longus (EDL) and soleus (SOL), which have distinctly different myosin heavy chain compositions (Burkholder et al., 1994), thus allowing us to address the potential role of muscle fiber type. Considering the literature, our hypothesis was that intrinsic skeletal muscle contractile function, fatigability and contraction-stimulated glucose uptake would be higher in the afternoon or evening, as compared to morning, regardless of the muscle studied.

## Material and Methods

### Animals

All studies were conducted in female C57BL/6NJ mice (The Jackson Laboratory, stock #05304) at 13.0 ± 0.1 weeks of age. Animals arrived at the vivarium at 10 weeks of age and were housed in a conventional facility with a 12-hour light/12-hour dark cycle (Light: 0600 h [ZT0]; Dark: 1800 h [ZT12]) for 3.0 ± 0.2 weeks after arrival.

There were four experimental groups: ZT1, ZT7, ZT13, ZT19; the experimental timing is overviewed in Figure 1A. These times during the light (ZT1 and ZT7) and dark (ZT13 and ZT19) phase were chosen to represent “early” (ZT1 and ZT13) and “late” (ZT7 and ZT19) timepoints within each phase. If the experiment occurred during a dark phase timepoint, all animal handling was done under dim red light. After anesthetization, muscle dissections and *ex vivo* testing occurred under normal ambient light. Because eating patterns differ by time-of day in mice and meal timing and carbohydrate intake impacts exercise performance (Aoyama and Shibata, 2020; Coyle et al., 1985; Kerksick et al., 2017), we controlled the last meal before tissue dissection. Thus, all mice were orally gavaged with 50% dextrose (2 g/kg) 3 hours before tissue dissection and were then fasted with *ad libitum* access to water. All animal experiments were approved by and conducted in accordance with the Animal Care Program at the University of California, San Diego; Institutional Animal Care and Use Committee (IACUC) protocol #S09322.

**Figure 1.**
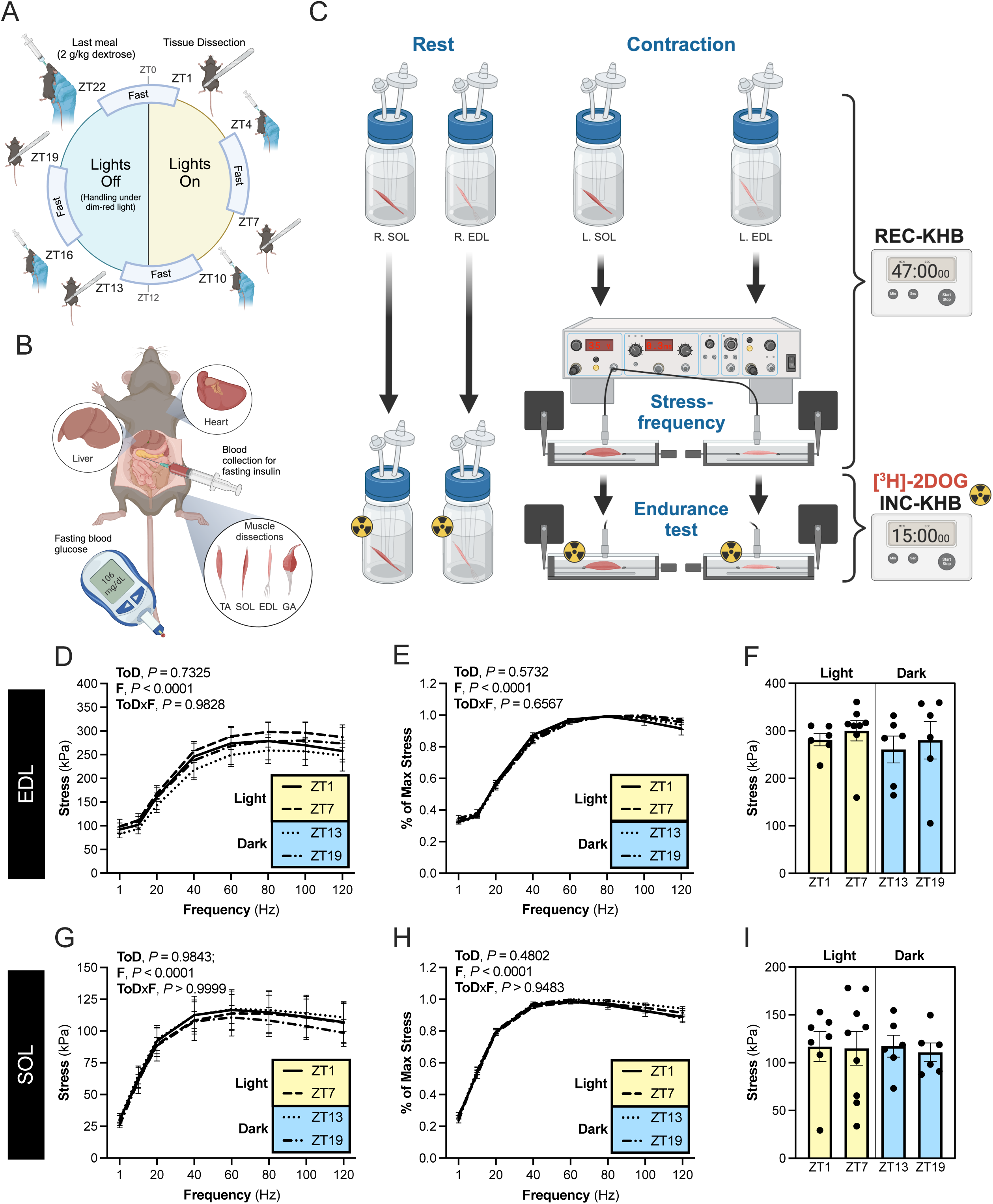
Intrinsic contractile function in two different skeletal muscles does not vary over a 24 h circadian cycle. **(A)** Overview of the experimental timing for the four experimental groups. Mice were orally gavaged with a standardized meal of 50% dextrose (2 g/kg) and then fasted for 3 hours before tissue dissection, which occurred at Zeitgeber Time (ZT) 1, ZT7, ZT13 and ZT19. Vivarium lights were on at ZT0 (0600 h) and were off at ZT12 1800 h). Thus, ZT1 and ZT7 were during the light/rest phase (yellow), whilst ZT13 and ZT19 were during the dark/active phase (blue). **(B)** Overview of the tissue dissection and collection procedure. TA, tibialis anterior; SOL, soleus; EDL, extensor digitorum longus; GA, gastrocnemius. **(C)** Overview of the *ex vivo* testing of intrinsic contractile function and endurance, and basal and contraction-stimulated [^3^H]-2-deoxyglucose (2DOG) uptake in paired (R., Right; L., Left) SOL and EDL. KHB, Krebs-Henseleit buffer; REC-KHB, recovery KHB; INC-KHB, incubation KHB. **(D)** Stress-frequency relationship in the EDL (1-120Hz, 300 ms train, 0.3 ms pulse, 35 V). **(E)** Stress-frequency relationship in the EDL, normalized to maximum tetanic stress. **(F)** Maximal tetanic stress in the EDL, extracted from the stress-frequency test. **(G)** Stress-frequency relationship in the SOL (1-120 Hz, 400 ms train, 0.3 ms pulse, 35 V). **(H)** Stress-frequency relationship in the SOL, normalized to maximum tetanic stress. **(I)** Maximal tetanic stress for the SOL, extracted from the stress-frequency testing. Data reported as mean±SEM. Statistics: **(D, E, G, H)** Repeated measures 2-way (ToD, time of day; F, stimulation frequency) ANOVA with Geisser-Greenhouse correction. **(F** and **I)** Ordinary one-way analysis of variance (ANOVA).

### Tissue dissection

An overview of tissue dissection is provided in Figure 1B. Fasted (3 h) mice were weighed (nearest 0.01g) and blood glucose (tail vein; Contour® blood glucose meter) was measured. Mice were then administered an intraperitoneal injection of a pentobarbital/phenytoin-containing solution (300 mg/kg; EUTHASOL®; Virbac, ANADA # 200-071). First, paired EDL and SOL were rapidly dissected and incubated in “recovery” Krebs-Henseleit buffer (REC-KHB; 116 mM NaCl, 4.6 mM KCl, 1.2 mM KH_2_PO_4_, 25 mM NaHCO_3_, 2.5 mM CaCl_2_, 1.2 mM MgSO_4_ at room temperature) for subsequent testing of contractile function and 2-deoxyglucose (2DOG) uptake (see, “*Ex-vivo SOL and EDL incubation: Assessment of contractile function and 2DOG uptake”* below). Blood was then collected from the inferior vena cava into EDTA-containing tubes, centrifuged (14,167 x g, 20 min, 4°C), and the plasma isolated.

Immediately after blood collection, tibialis anterior (TA), gastrocnemius (GA), liver and heart were dissected, weighed and rapidly frozen in liquid nitrogen. In a separate cohort of identically treated mice, we collected EDL and SOL for gene expression analysis, with these muscles being rapidly frozen (liquid nitrogen) after dissection. All samples were stored at -80°C.

### Ex-vivo EDL and SOL incubation: Assessment of contractile function and 2DOG uptake

The experimental design is overviewed in Figure 1C. Immediately after dissection, each EDL and SOL recovered in room temperature (21.2 ± 0.2 °C) REC- KHB, with each muscle incubated in a separate, oxygenated (95% O_2_, 5% CO_2_) flask. After 20 min, the SOL and EDL for contraction (i.e. CXN group) were transferred to a custom chamber for mechanical testing and assessment of glucose uptake. Thus, the muscle origin was tied with 4-0 silk suture to a rigid post, and the insertion was secured to the arm of a dual-mode ergometer. Then, muscles were electrically stimulated (model S88; Astro-Med, West Warwick, RI, USA) via parallel platinum electrodes (35 V, 0.3 ms pulse duration) with single twitches (1 Hz) to set optimal muscle length. After, 30 minutes in REC-KHB, the CXN group EDL and SOL underwent a two-step contraction protocol; the first step was to test the stress-frequency relationship and the second step was to test fatigability. The stress-frequency testing was as follows: muscles were stimulated (EDL: 300 ms train, 0.3 ms pulse; SOL: 400 ms train, 0.3 ms pulse) at difference frequencies (1, 10, 20, 40, 60, 80, 100, 120 Hz), with 60 seconds between each contraction. After assessing stress-frequency, the muscles rested for 10 minutes, with the total time to complete the stress-frequency curve and rest being 17 min. Then, the REC-KHB was replaced with room temperature “incubation” KHB (INC-KHB: 116 mM NaCl, 4.6 mM KCl, 1.2 mM KH_2_PO_4_, 25 mM NaHCO_3_, 2.5 mM CaCl_2_, 1.2 mM MgSO_4_, 2 mM Na-pyruvate, 8 mM mannitol, 1 mM 2DOG, 9 mM [^14^C]-mannitol [55 mCi/mmol; American Radiolabeled Chemicals, Inc.], and 1 mM [^3^H]-2-deoxyglucose [2DOG] [6 mCi/mmol; American Radiolabeled Chemicals, Inc.]) for the assessment of 2DOG uptake during contraction. Thus, immediately after switching to INC-KHB, each muscle underwent a fatiguing contraction protocol for 15 min (EDL: 1,000 ms train, 0.3 ms pulse, 100 Hz every 15 sec; SOL: 400 ms train, 0.3 ms pulse, 40 Hz every 2 sec). For the contralateral SOL and EDL, which were in the “rested” group (Rest), after a comparable 47 minutes of incubation in REC-KHB, the EDL and SOL were transferred to flasks containing room temperature INC-KHB for the assessment of basal 2DOG uptake. After 15 min in INC-KHB, all CXN and Rest muscles were rapidly blotted dry, trimmed, weighed, frozen in liquid nitrogen, and stored (-80°C). Muscle stresses were calculated by normalizing force in Newtons to the physiologic cross-sectional area (PCSA) of each muscle (Stress [kPa] = F [Newtons]/PCSA [mm^2^]). Cumulative fatigue was measured by assessing area under the curve (units = kPa•min).

### Muscle homogenization for 2DOG uptake assessment and immunoblotting

Muscles were transferred to 1.5 mL tubes on ice containing a 1/8 inch stainless steel bead and 500 μL of homogenization buffer (50 mm Tris [pH 7.5], 250 mM sucrose, 1 mM EDTA, 1 mM EGTA, 1% Triton X-100, 50 mM NaF, 1 mM NaCO_2_Na_2_(PO_4_)_2_, and 0.1% DTT) containing 1 M nicotinamide (MilliporeSigma #N0636), 1 mM Pefabloc SC PLUS (MilliporeSigma #11873601001), 1 mM trichostatin A (Cell Signaling #9950S), Complete (MilliporeSigma #11836170001), phosphatase inhibitor cocktail (PIC) 2 (MilliporeSigma #P5726), and PIC3 (MilliporeSigma #P0044). The muscles were then homogenized (Bullet Blender, Next Advance #BT24M) and subsequently rotated for 2 hours at 4°C. The homogenate was then centrifuged (14,489 x g) for 20 minutes at 4°C. The supernatant was collected and stored at -80° C for subsequent scintillation counting and determination of 2DOG uptake as previously described (McCurdy and Cartee, 2005). Immunoblotting was conducted using the Jess Automated Western Blot System (Protein Simple #004-650). Antibodies used were phospho-AMPKα (Thr172; pAMPK [T172]) Antibody (Cell Signaling #2531) and eEF2 (Cell Signaling #2332).

### RNA extraction, reverse transcription, and real-time PCR

RNA was extracted from tissues with TRIzol Reagent (Invitrogen™ #15596026). RNA concentration and quality of RNA was measured (NanoDrop™ 2000 spectrophotometer; Thermo Scientific™ #ND-2000) and 500 ng of RNA was used for cDNA synthesis (Applied Biosystems, #4368814). Semi**-**quantitative real-time PCR analysis was conducted using PowerUp™ SYBR™ Green master mix (Thermo Scientific™ #A25741). Relative expression levels for each gene of interest were calculated with the ΔΔCt method, using *Rn18s* as the normalization control. Primers used were: *Bmal1* (5’- CACTGTCCCAGGCATTCCA-3’ FWD, 5’- TTCCTCCGCGATCATTCG-3’ REV), *Dbp* (5’- CCTGAGGAACAGAAGGATGA-3’ FWD, 5’-ATCTGGTTCTCCTTGAGTCTT-3’ REV), *Nr1d1* (5’- TGGCCTCAGGCTTCCACTATG-3’ FWD, 5’- CCGTTGCTTCTCTCTCTTGGG-3’ REV), *Rn18s* (5’- GCTTAATTTGACTCAACACGGGA-3’ FWD, 5’- AGCTATCAATCTGTCAATCCTGTC-3’ REV).

### Plasma insulin

Plasma insulin was analyzed using an ELISA kit, per the manufacturer’s instructions (80-INSMS-E-01; ALPCO Diagnostics).

### Statistics

Statistical analyses were performed using Prism v10.0.2 (GraphPad Software Inc., La Jolla, CA, USA). For Figure 1 and 2, data were analyzed by either a 1- or 2-way analysis of variance or mixed-effects model; details regarding specific statistical tests for each figure are detailed in corresponding figure legends. For Figure 3, statistical significance of circadian rhythmicity was determined by a zero-amplitude test on the Cosinor.Online web application (Molcan, 2023). All data are expressed as mean±SEM. Significant differences (*P* < 0.05) are marked with “#” in figures. Sample sizes to detect a 20% difference with an a of 0.05 and a b of 80% were estimated using G*Power v3.1.9.6 and were based on means and standard deviations from the literature for maximal tetanic tension (Svensson et al., 2020), fatigability (Jensen et al., 2014), and contraction-stimulated glucose uptake (Kang et al., 2021).

**Figure 2.**
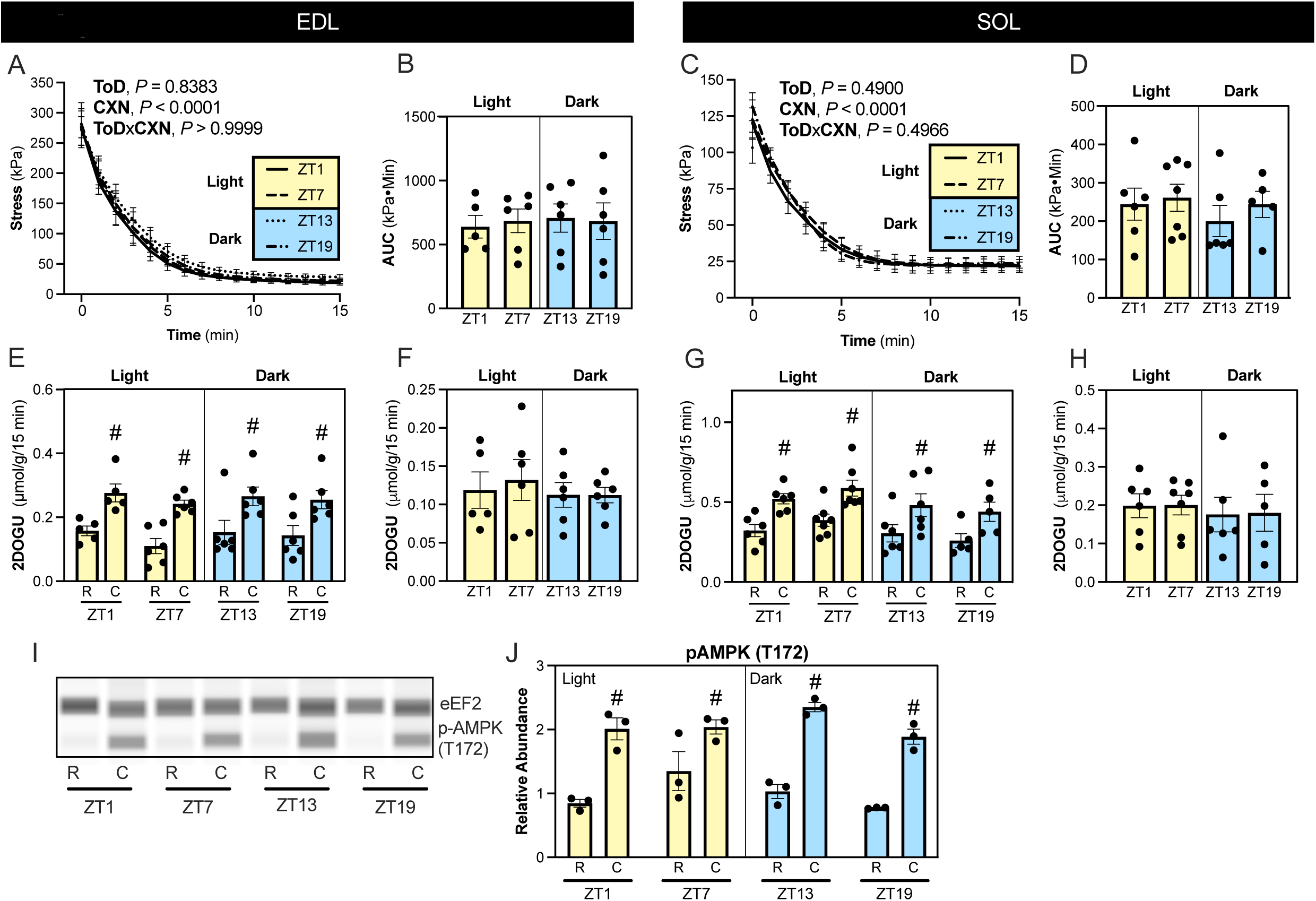
Muscle endurance and contraction-stimulated glucose uptake does not vary over a 24 h circadian cycle. All measurements for all panels were conducted at ZT1, ZT7, ZT13, and ZT19. **(A)** Muscle stresses measured during the fatigue test for EDL. **(B)** Cumulative fatigue during the endurance test, measured by AUC (kPa•Min), for EDL. **(C)** Muscle stresses measured during the fatigue test for SOL. **(D)** Cumulative fatigue during the endurance test, measured by AUC (kPa•Min), for SOL. **(E)** 2DOG uptake in rested (R) and contracted (C) EDL. **(F)** Contraction-stimulated glucose uptake (Contraction 2DOG uptake – Rested 2DOG uptake) in EDL. **(G)** 2DOG uptake in rested (R) and contracted (C) SOL. **(H)** Contraction-stimulated glucose uptake in SOL. **(I)** Representative immunoblot for eEF2 and pAMPK (T172) in rested (R) and contracted (C) EDL. **(J)** Relative abundance of pAMPK (T172) in rested (R) vs. contracted (C) EDL (normalized to eEF2). Data reported as mean±SEM. Statistics: **(A** and **C)** Repeated measures 2way ANOVA with Geisser-Greenhouse correction, Tukey’s multiple comparisons test. **(B**, **D**, **F**, and **H)** Ordinary one-way ANOVA with Tukey’s multiple comparison’s test. **(E, G,** and **J)** Repeated measures 2-way ANOVA with Šídák’s multiple comparison’s test.

**Figure 3.**
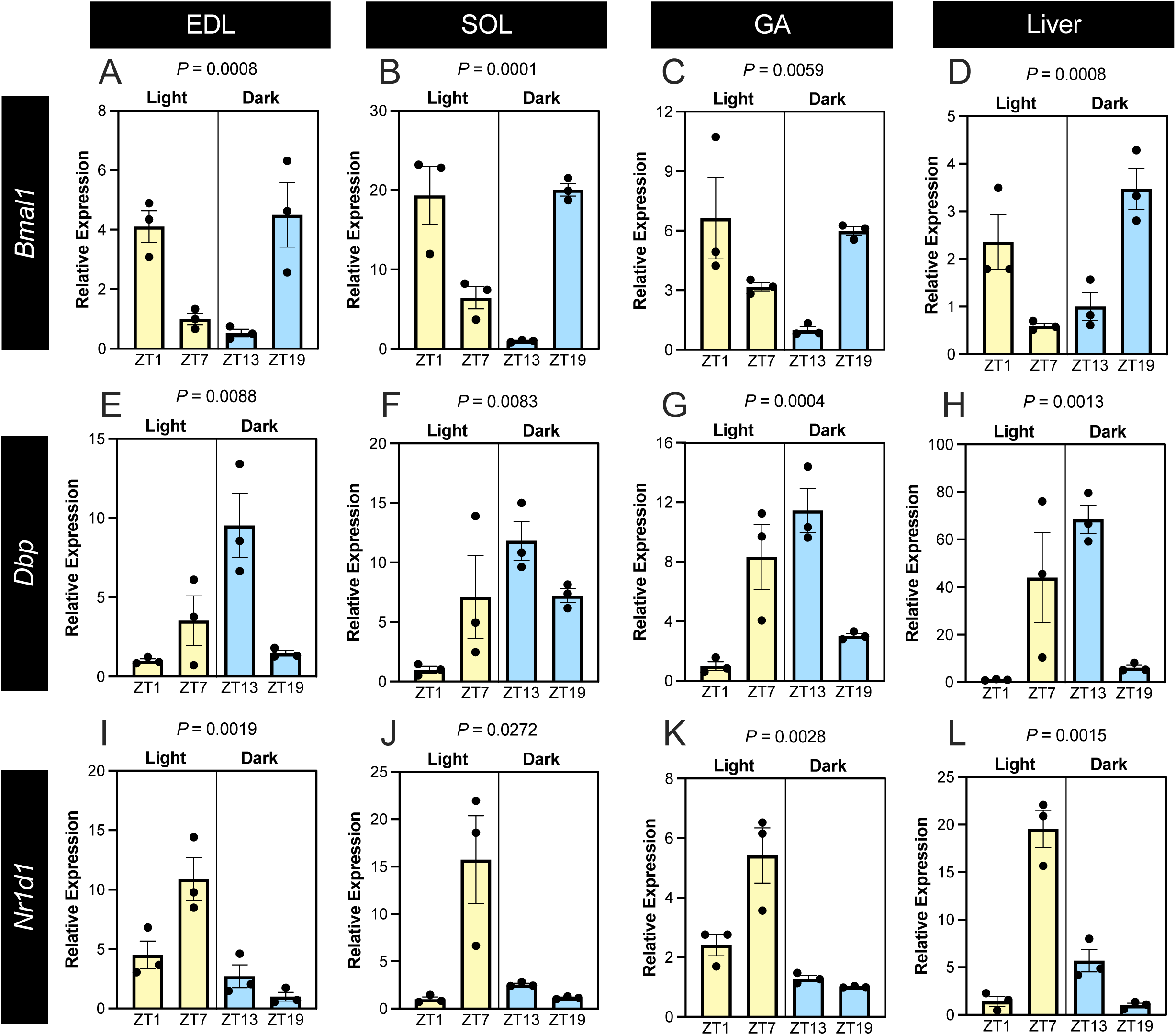
Normal circadian rhythmicity in the expression of core clock genes in skeletal muscle and liver. All measurements for all panels were conducted in tissues collected at ZT1, ZT7, ZT13, and ZT19. mRNA expression of *Bmal1* **(A-D)**, *Dbp* **(E-H)**, and *Nr1d1* **(I-L)** normalized to *Rn18s* expression in EDL **(A, E,** and **I)**, SOL **(B, F,** and **J)**, gastrocnemius [GA **(C, G,** and **K)**], and liver **(D, H,** and **L)**. Data reported as mean±SEM. Statistics: Zero amplitude test for circadian rhythmicity (*P* reported for each gene-tissue combination above each panel).

## Results and Discussion

### Intrinsic skeletal muscle contractile function does not vary over a 24 h circadian cycle

Studies suggest that circadian variation in skeletal muscle contractile function, such as anaerobic power output, concentric force production, and maximal torque production, is due to diurnal variation in the intrinsic contractile properties of the skeletal muscle (Chtourou et al., 2011; Douglas et al., 2021; Guette et al., 2005; Martin et al., 1999b; Nicolas et al., 2007). Others suggest that daily variance in exercise capacity in mice may be due to distinct diurnal transcriptomic and metabolic signatures (e.g. NAD^+^, ZMP) in skeletal muscle (Ezagouri et al., 2019; McCarthy et al., 2007; Peek et al., 2013). Certainly, the time-of-day that exercise is undertaken has differential effects on the gene expression response to exercise (Casanova-Vallve et al., 2022; Ezagouri et al., 2019; Maier et al., 2022; Mirizio et al., 2020). However, a key gap in the field is that, to our knowledge, no studies have investigated skeletal muscle contractile function at multiple points throughout both the light and dark phases of the circadian cycle (i.e., over a 24 h period).

For our study, body mass, muscle mass (TA, SOL, and EDL), liver and heart mass, and fasting glucose and insulin concertation were not different across the 4 timepoints (Supplementary Table 1). As it relates to intrinsic force generating capacity, for both EDL and SOL, while there was the expected effect of stimulation frequency on muscle stress production, contrary to our hypothesis and current thinking in the field, the stress-frequency relationship at low, moderate, or maximal stimulation frequency was not different across a 24 h circadian cycle, in either muscle (EDL: Figure 1D and 1F, Supplementary Table 1 [twitch parameters] and Supplementary Figure 1A-D; SOL: Figure 1G and 1I, Supplementary Table 1 [twitch parameters] and Supplementary Figure 1E-H). Notably, while Kahn et al., (2024) did not study contractile function across the range of stimulation frequencies, our findings of no difference in maximal tetanic stress in the EDL during the light phase is in line with their maximal tetanic stress data (personal communication with Dayanidhi S, Kahn RE, Lieber RL). There was also no time-of-day effect in the EDL or SOL when expressing the stress frequency curve relative to maximum stress (Figure 1E and 1H, respectively). Overall, by measuring muscle intrinsic contractile properties at multiple time points across both the light and dark phases of the circadian cycle, and across a wide range of stimulation frequencies, these data demonstrate that the intrinsic contractile properties of skeletal muscle do not vary over the course of a 24 h circadian cycle in female mice, regardless of muscle myosin heavy chain composition.

### Muscle endurance does not vary over a 24 h circadian cycle

Similar to the effects of time-of-day on muscle force production, there is a documented effect of time-of-day on endurance exercise performance across species, including in humans (Chtourou et al., 2011; Gemmink et al., 2023; Souissi et al., 2002; van Moorsel et al., 2016) and rodents (Adamovich et al., 2021; Ezagouri et al., 2019; Maier et al., 2022; Wolff & Esser, 2012). A potential role of the circadian clock in endurance performance is further substantiated by the fact that mouse models with disruption of the circadian clock (e.g. *Per2^-/-^* and *Bmal1^-/-^* mice) do not demonstrate variability in exercise performance within and/or between the light and dark phases of the circadian cycle (Adamovich et al., 2021; Ezagouri et al., 2019; Hesketh et al., 2023; Xin et al., 2023).

Nevertheless, while some studies demonstrate that treadmill running performance in mice differs between the light and dark phases in wild-type mice (Ezagouri et al., 2019; Maier et al., 2022), importantly, this is not a universal finding (Casanova-Vallve et al., 2022). While, variability in these results could be due to many factors, including the treadmill testing approach used (e.g. electrical shock or not), as well as other common factors impacted by time-of-day, such as body temperature, motivation to exercise, food intake, muscle glycogen content, central nervous system arousal, and pain tolerance, intrinsic muscle properties have been emphasized as an underlying mediator (Douglas et al., 2021), although this concept has not been empirically tested.

Thus, to specifically study intrinsic skeletal muscle endurance over a 24 h circadian cycle, we conducted a fatiguing contraction protocol in the EDL and SOL. To confirm that the muscles were not fatigued before starting the fatiguing protocol, we compared the stresses for the first contraction of the fatigue protocol to the corresponding stimulation frequency from the stress-frequency curve. Importantly, there was no significant difference in the stresses at these time points, in either EDL or SOL (Supplementary Figure 2A and 2B, respectively). During the endurance protocol, as expected, muscle stresses robustly decreased in response to repeated contractions in the EDL (Figure 2A and 2B) and SOL (Figure 2C and 2D). Significantly however, the rate of fatigability did not differ within the light or dark phases, or between the light and dark phases, in either muscle.

As expected, pAMPK (T172) was significantly increased by the protocol, but there was no time-of-day variability within the resting or contracting groups (Figure 2J). This lack of an effect of time-of-day on basal or exercise-mediated activation of pAMPK (T172) in skeletal muscle is in line with work by others (Ezagouri et al., 2019), but differs from findings in murine hypothalamus and embryonic fibroblasts, which demonstrate circadian oscillations in pAMPK (T172) (Um et al., 2011). In summary, contrary to our hypothesis and current thinking in the field, intrinsic skeletal muscle endurance does not vary over the course of a 24 hour circadian cycle in female mice, regardless of muscle type.

### Contraction-stimulated glucose uptake by skeletal muscle does not vary by time-of-day

To meet the energetic demands of contraction, skeletal muscle glucose uptake increases over time and/or with increasing intensity of contraction (Fell et al., 1982; Ploug et al., 1984b; Richter and Hargreaves, 2013; Romijn et al., 1993). In addition to meeting the energetic demands of exercise, this contraction-stimulated glucose uptake is important to glycemic control, with exercise being a cornerstone intervention for treating or preventing clinical hyperglycemia (“Reduction in the Incidence of Type 2 Diabetes with Lifestyle Intervention or Metformin,” 2002).

Nevertheless, while studies (Mancilla et al., 2021; Qian et al., 2023; Savikj et al., 2019) demonstrate time-of-day effects of exercise training on glycemic control, we studied whether this might be due to intrinsic changes contraction-stimulated glucose uptake by muscle. As expected, there was a robust effect of contraction to increase muscle glucose uptake as compared to the contralateral rested muscle in the EDL (∼85% higher in CXN vs. Rest; Figure 2E) and SOL (∼59% higher in CXN vs. Rest; Figure 2G). However, there were no time-of-day difference in 2DOG uptake when comparing within the rested or contracted muscles. As a result, contraction-stimulated 2DOG uptake (calculated as: CXN 2DOG uptake – Rest 2DOG uptake) was not different across the 24 h circadian cycle in the EDL or SOL (Figure 2F and 2H, respectively). Thus, contrary to current thinking in the field, contraction-stimulated glucose uptake does not vary over the course of a 24 h circadian cycle in female mice, regardless of muscle type.

### Normal circadian rhythmicity in gene expression in skeletal muscle and liver

Food intake and carbohydrate content of meals are important contributors to exercise performance (Anantaraman et al., 1995; Costill et al., 1981; Coyle et al., 1985; Fielding et al., 1985; Kerksick et al., 2017) and a powerful zeitgeber for the circadian cycle (Aoyama and Shibata, 2020; Stephan, 2002). Moreover, systemic glucose or insulin (through carbohydrate intake) availability can profoundly impact muscle gene expression (Batista et al., 2019). To date, to our knowledge, all studies that have focused on the role of time-of-day on exercise performance have not controlled for the potential effect of food/carbohydrate availability, which may underlie some of the variability in findings in the field. Addressing this, in our study all mice received a standardized glucose meal 3 hours before tissue dissection. Accordingly, we wanted to validate that there was still a circadian rhythmicity in skeletal muscle and liver (as an example of another important metabolic tissue). Thus, we measured mRNA expression levels of the core clock regulator *Bmal1,* and the clock output genes *Dbp* and *Nr1d1* in EDL (Figure 3A, E, and I, respectively), SOL (Figure 3B, F, and J, respectively), GA (Figure 3C, G, and K, respectively), and liver (Figure 3D, H, and L, respectively). In these three muscles and the liver, there was significant circadian rhythmicity of the three genes (as measured by zero-amplitude testing), with circadian changes in these genes being consistent with previous studies in skeletal muscle (McCarthy et al., 2007) and liver (Guan and Lazar, 2022). These data demonstrate that circadian rhythmicity, regardless of prior food intake, is strongly entrained in skeletal muscle and liver. It also demonstrates that the lack of effect of time-of-day on skeletal muscle contractile function, fatigability or contraction-stimulated glucose uptake is not due to a lack of circadian rhythmicity.

### Summary

Contrary to our hypothesis and current thinking in the field, we found that the intrinsic functionality of two distinctly different skeletal muscles, as well as contraction-stimulated glucose uptake by these muscles, does not differ over a 24 h circadian cycle in female mice. Thus, circadian variation in exercise performance and the glycemia-reducing benefits of exercise is not due to chronobiological effects on intrinsic muscle function or contraction-stimulated glucose uptake, respectively. In future work, it will be of interest to validate these findings in male mice, and to further refine the underlying factor(s) (e.g. body [or muscle] temperature, meal timing, muscle glycogen, motor unit recruitment, humoral factors, etc.) that mediate chronobiological variation in exercise performance or how exercise modulates systemic glycemia.

## Acknowledgements

The authors thank Dr. Sudarshan Dayanidhi, Dr. Richard L. Lieber, and Ryan E. Kahn for insightful and thoughtful personal communications on their study on the effect of time-of-day on *ex vivo* muscle contractility following short-term satellite cell ablation. This work was supported, in part, by National Institutes of Health (NIH) grants R21 AR072882 and R21 AG067495 to S. Schenk, whilst L.S. Fitzgerald was supported, in part, by the NIH-funded UC San Diego Medical Scientist Training Program (T32 GM007198) and a Summer Research Fellowship from the UC San Diego School of Medicine. The authors also acknowledge support from the Wu-Tsai Human Performance Alliance and the Joe and Clara Tsai Foundation.

## Author Details

Liam S. Fitzgerald; conception and design, acquisition of data, analysis and interpretation of data, drafting and revising the article. Shannon N. Bremner; acquisition, analysis and interpretation of data, revising the article. Samuel R. Ward; analysis and interpretation of data, revising the article and funding acquisition. Yoshitake Cho: acquisition of data, analysis and interpretation of data, revising the article. Simon Schenk; conception and design, acquisition of data, analysis and interpretation of data, drafting and revising the article and funding acquisition.

## Funding

**National Institute on Aging (R21AG067495).** Simon Schenk.

**National Institute of Arthritis and Musculoskeletal and Skin Diseases (R21AR072882).** Simon Schenk.

**National Institute of General Medical Sciences (T32GM007198).** Liam S. Fitzgerald; through the MSTP program within the School of Medicine at the University of California, San Diego.

**School of Medicine at the University of California, San Diego (Summer Research Fellowship).** Liam S. Fitzgerald.

**Wu-Tsai Human Performance Alliance and the Joe and Clara Tsai Foundation.** Samuel R. Ward, Simon Schenk.

---

**Supplementary Figure 1. Force tracings for EDL and SOL at different times-of-day.** Example force tracings for each stimulation frequency (i.e. 1, 10, 20, 40, 60, 80, 100, 120 Hz) during the stress-frequency testing in the **(A-D)** EDL (300 ms train, 0.3 ms pulse, 35 V), and **(E-H)** SOL (400 ms train, 0.3 ms pulse, 35 V) at ZT1, ZT7, ZT13, and ZT19.

**Supplementary Figure 2. Comparison of tetanic tension during the stress-frequency testing versus the initial contraction of the endurance test.** Comparison of forces at the same stimulation frequency from the stress-frequency test (SF) versus the first contraction of the fatigue test (FT) in the **(A)** EDL (100 Hz, 300 ms train, 0.3 ms pulse) and **(B)** SOL (40 Hz, 400 ms train, 0.3 ms pulse) at ZT1, ZT7, ZT13, and ZT19. Data reported as mean±SEM. Statistics: unpaired t-test within each time point.

**Supplementary Table 1.** Body and tissue masses, fasting glucose and insulin, and contractile twitch characteristics at different times-of-day. Abbreviations: EDL, extensor digitorum longus; SOL, soleus; TA, tibialis anterior; ZT, zeitgeber time; PTT, peak twitch tension; TTP, time-to-peak twitch tension. Statistics: Within each row, data were analyzed by one-way ANOVA with Tukey’s multiple comparison’s test. Data are expressed as mean±SEM.

|  | **ZT1** | **ZT7** | **ZT13** | **ZT19** | **P**<br>**value** | **n**<br>**(ZT1/7/13/19)** |
| --- | --- | --- | --- | --- | --- | --- |
| **Body and tissue mass** |  |  |  |  |  |  |
| **Body (g)** | 26.7±0.6 | 25.8±0.9 | 25.6±0.4 | 25.6±0.6 | 0.9587 | 7/9/8/6 |
| **EDL (mg)** | 11.5±0.6 | 11.0±0.7 | 12.2±0.4 | 12.0±0.5 | 0.3432 | 7/9/8/6 |
| **SOL (mg)** | 11.3±0.5 | 11.2±0.6 | 11.8±0.5 | 11.5±0.3 | 0.6616 | 7/9/8/6 |
| **TA (mg)** | 51.6±1.1 | 50.3±1.7 | 51.4±1.6 | 49.5±1.4 | 0.7709 | 7/9/8/6 |
| **Heart (mg)** | 121.1±3.2 | 124.0±6.0 | 128.3±4.0 | 113.7±5.0 | 0.1995 | 7/9/8/6 |
| **Liver (g)** | 1.32±0.06 | 1.20±0.04 | 1.18±0.06 | 1.17±0.06 | 0.9268 | 7/9/8/6 |
| **Fasting glucose and insulin** |  |  |  |  |  |  |
| **Fasting**<br>**glucose**<br>**(mg/dL)** | 131±6 | 119±5 | 109±5 | 112±8 | 0.4027 | 7/9/8/6 |
| **Fasting**<br>**insulin**<br>**(ng/mL)** | 0.85±0.04 | 0.88±0.06 | 0.82±0.06 | 0.83±0.06 | 0.7686 | 4/4/4/4 |

| Contractile function: Twitch characteristics |  |  |  |  |  |  |  |
| --- | --- | --- | --- | --- | --- | --- | --- |
| EDL | PTT<br>(kPa) | 92.0±5.4 | 98.7±7.2 | 83.0±8.6 | 98.7±15.1 | 0.6229 | 6/8/6/6 |
|  | TPT<br>(ms) | 25.3±0.6 | 24.4±0.8 | 25.3±0.9 | 23.7±0.9 | 0.9691 | 6/8/6/6 |
| SOL | PTT<br>(kPa) | 29.6±3.8 | 30.5±4.8 | 30.5±3.0 | 26.0±2.2 | 0.8507 | 7/9/6/6 |
|  | TTP<br>(ms) | 60.0±1.6 | 55.8±1.8 | 62.8±2.9 | 64.3±6.6 | 0.2905 | 7/9/6/6 |

## References

Adamovich Y, Dandavate V, Ezagouri S, Manella G, Zwighaft Z, Sobel J, Kuperman Y, Golik M, Auerbach A, Itkin M, Malitsky S, Asher G. 2021. Clock proteins and training modify exercise capacity in a daytime-dependent manner. Proc Natl Acad Sci U S A 118. doi:10.1073/PNAS.2101115118/

Anantaraman R, Carmines AA, Gaesser GA, Weltman A. 1995. Effects of carbohydrate supplementation on performance during 1 hour of high-intensity exercise. Int J Sports Med 16:461–465. doi:10.1055/s-2007-973038

Aoyama S, Shibata S. 2020. Time-of-Day-Dependent Physiological Responses to Meal and Exercise. Front Nutr 7:18. doi:10.3389/FNUT.2020.00018

Atkinson G, Reilly T. 1996. Circadian variation in sports performance. Sports Med 21:292–312. doi:10.2165/00007256-199621040-00005

Atkinson G, Speirs L. 1998. Diurnal variation in tennis service. Percept Mot Skills 86:1335–1338. doi:10.2466/PMS.1998.86.3C.1335

Bajpeyi S, Tanner CJ, Slentz CA, Duscha BD, McCartney JS, Hickner RC, Kraus WE, Houmard JA. 2009. Effect of exercise intensity and volume on persistence of insulin sensitivity during training cessation. J Appl Physiol (1985) **106**:1079–1085. doi:10.1152/JAPPLPHYSIOL.91262.2008

Balducci S, Iacobellis G, Parisi L, Di Biase N, Calandriello E, Leonetti F, Fallucca F. 2006. Exercise training can modify the natural history of diabetic peripheral neuropathy. J Diabetes Complications 20:216–223. doi:10.1016/J.JDIACOMP.2005.07.005

Batista TM, Garcia-Martin R, Cai W, Konishi M, O’Neill BT, Sakaguchi M, Kim JH, Jung DY, Kim JK, Kahn CR. 2019. Multi-dimensional Transcriptional Remodeling by Physiological Insulin In Vivo. Cell Rep 26:3429–3443.e3. doi:10.1016/J.CELREP.2019.02.081

Baxter C, Reilly T. 1983. Influence of time of day on all-out swimming. Br J Sports Med 17:122. doi:10.1136/BJSM.17.2.122

Bergh U, Ekblom B. 1979. Influence of muscle temperature on maximal muscle strength and power output in human skeletal muscles. Acta Physiol Scand 107:33–37. doi:10.1111/J.1748-1716.1979.TB06439.X

Bessot N, Nicolas A, Moussay S, Gauthier A, Sesboüé B, Davenne D. 2006. The effect of pedal rate and time of day on the time to exhaustion from high-intensity exercise. Chronobiol Int 23:1009–1024. doi:10.1080/07420520600920726

Burkholder TJ, Fingado B, Baron S, Lieber RL. 1994. Relationship between muscle fiber types and sizes and muscle architectural properties in the mouse hindlimb. J Morphol 221:177–190. doi:10.1002/JMOR.1052210207

Casanova-Vallve N, Duglan D, Vaughan ME, Pariollaud M, Handzlik MK, Fan W, Yu RT, Liddle C, Downes M, Delezie J, Mello R, Chan AB, Westermark PO, Metallo CM, Evans RM, Lamia KA. 2022. Daily running enhances molecular and physiological circadian rhythms in skeletal muscle. Mol Metab 61. doi:10.1016/J.MOLMET.2022.101504

Chtourou H, Zarrouk N, Chaouachi A, Dogui M, Behm DG, Chamari K, Hug F, Souissi N. 2011. Diurnal variation in Wingate-test performance and associated electromyographic parameters. Chronobiol Int 28:706–713. doi:10.3109/07420528.2011.596295

Cohen ND, Dunstan DW, Robinson C, Vulikh E, Zimmet PZ, Shaw JE. 2008. Improved endothelial function following a 14-month resistance exercise training program in adults with type 2 diabetes. Diabetes Res Clin Pract 79:405–411. doi:10.1016/J.DIABRES.2007.09.020

Constable SH, Favier RJ, Cartee GD, Young DA, Holloszy JO. 1988. Muscle glucose transport: interactions of in vitro contractions, insulin, and exercise. 101152/jappl19886462329 **64**:2329–2332. doi:10.1152/JAPPL.1988.64.6.2329

Costill DL, Sherman WM, Fink WJ, Maresh C, Witten M, Miller JM. 1981. The role of dietary carbohydrates in muscle glycogen resynthesis after strenuous running. American Journal of Clinical Nutrition 34:1831–1836. doi:10.1093/ajcn/34.9.1831

Coyle EF, Coggan AR, Hemmert MK, Lowe RC, Walters TJ. 1985. Substrate usage during prolonged exercise following a pre-exercise meal. J Appl Physiol 59:429–433. doi:10.1152/jappl.1985.59.2.429

Czelusniak O, Favreau E, Ives SJ. 2021. Effects of Warm-Up on Sprint Swimming Performance, Rating of Perceived Exertion, and Blood Lactate Concentration: A Systematic Review. J Funct Morphol Kinesiol 6. doi:10.3390/JFMK6040085

Douglas CM, Hesketh SJ, Esser KA. 2021. Time of day and muscle strength: A circadian output? Physiology 36:44–51. doi:10.1152/PHYSIOL.00030.2020/

Edwards BJ, Lindsay K, Waterhouse J. 2005. Effect of time of day on the accuracy and consistency of the badminton serve. Ergonomics 48:1488–1498. doi:10.1080/00140130500100975

Evans EM, Racette SB, Peterson LE, Villareal DT, Greiwe JS, Holloszy JO. 2005. Aerobic power and insulin action improve in response to endurance exercise training in healthy 77-87 yr olds. J Appl Physiol (1985) **98**:40–45. doi:10.1152/JAPPLPHYSIOL.00928.2004

Ezagouri S, Zwighaft Z, Sobel J, Baillieul S, Doutreleau S, Ladeuix B, Golik M, Verges S, Asher G. 2019. Physiological and Molecular Dissection of Daily Variance in Exercise Capacity. Cell Metab 30:78–91.e4. doi:10.1016/j.cmet.2019.03.012

Fell RD, Terblanche SE, Ivy JL, Young JC, Holloszy JO. 1982. Effect of muscle glycogen content on glucose uptake following exercise. J Appl Physiol Respir Environ Exerc Physiol 52:434–437. doi:10.1152/JAPPL.1982.52.2.434

Fielding RA, Costill DL, Fink WJ, King DS, Hargreaves M, Kovaleski JE. 1985. Effect of carbohydrate feeding frequencies and dosage on muscle glycogen use during exercise. Med Sci Sports Exerc 17:472–476. doi:10.1249/00005768-198508000-00012

Gemmink A, Daemen S, Wefers J, Hansen J, van Moorsel D, Astuti P, Jorgensen JA, Kornips E, Schaart G, Hoeks J, Schrauwen P, Hesselink MKC. 2023. Twenty-four hour rhythmicity in mitochondrial network connectivity and mitochondrial respiration; a study in human skeletal muscle biopsies of young lean and older individuals with obesity. Mol Metab 72. doi:10.1016/J.MOLMET.2023.101727

Goodyear LJ, Kahn BB. 1998. Exercise, glucose transport, and insulin sensitivity. Annu Rev Med 49:235–261. doi:10.1146/ANNUREV.MED.49.1.235

Goodyear LJ, King PA, Hirshman MF, Thompson CM, Horton ED, Horton ES. 1990. Contractile activity increases plasma membrane glucose transporters in absence of insulin. Am J Physiol 258. doi:10.1152/AJPENDO.1990.258.4.E667

Guan D, Lazar MA. 2022. Circadian regulation of gene expression and metabolism in the liver. Semin Liver Dis 42:113. doi:10.1055/A-1792-4240

Gueldich H, Zarrouk N, Chtourou H, Zghal F, Sahli S, Rebai H. 2017. Electrostimulation Training Effects on diurnal Fluctuations of Neuromuscular Performance. Int J Sports Med 38:41–47. doi:10.1055/S-0042-115033

Guette M, Gondin J, Martin A. 2005. Time-of-day effect on the torque and neuromuscular properties of dominant and non-dominant quadriceps femoris. Chronobiol Int 22:541–558. doi:10.1081/CBI-200062407

Harrison SM, Bers DM. 1989. Influence of temperature on the calcium sensitivity of the myofilaments of skinned ventricular muscle from the rabbit. J Gen Physiol 93:411–428. doi:10.1085/JGP.93.3.411

Hesketh SJ, Sexton CL, Wolff CA, Viggars MR, Esser KA. 2023. Early morning run-training results in enhanced endurance performance adaptations in mice. bioRxiv. doi:10.1101/2023.09.18.557933

Jensen TE, Sylow L, Rose AJ, Madsen AB, Angin Y, Maarbjerg SJ, Richter EA. 2014. Contraction-stimulated glucose transport in muscle is controlled by AMPK and mechanical stress but not sarcoplasmatic reticulum Ca2+ release. Mol Metab 3:742–753. doi:10.1016/J.MOLMET.2014.07.005

Kahn RE, Lieber RL, Meza G, Dinnunhan F, Lacham-Kaplan O, Dayanidhi S, Hawley JA. 2024. Time-of-day effects on *ex vivo* muscle contractility following short-term satellite cell ablation. American Journal of Physiology-Cell Physiology. doi:10.1152/AJPCELL.00157.2024

Kang JH, Park JE, Dagoon J, Masson SWC, Merry TL, Bremner SN, Dent JR, Schenk S. 2021. Sirtuin 1 is not required for contraction-stimulated glucose uptake in mouse skeletal muscle. J Appl Physiol 130:1893. doi:10.1152/JAPPLPHYSIOL.00065.2021

Kerksick CM, Arent S, Schoenfeld BJ, Stout JR, Campbell B, Wilborn CD, Taylor L, Kalman D, Smith-Ryan AE, Kreider RB, Willoughby D, Arciero PJ, VanDusseldorp TA, Ormsbee MJ, Wildman R, Greenwood M, Ziegenfuss TN, Aragon AA, Antonio J. 2017. International society of sports nutrition position stand: nutrient timing. J Int Soc Sports Nutr 14:33. doi:10.1186/S12970-017-0189-4

Koch CE, Begemann K, Kiehn JT, Griewahn L, Mauer J, M. E. Hess, Moser A, Schmid SM, Brüning JC, Oster H. 2020. Circadian regulation of hedonic appetite in mice by clocks in dopaminergic neurons of the VTA. Nature Communications 2020 11:1 11:1–11. doi:10.1038/s41467-020-16882-6

Krauchi K, Wirz-Justice A. 1994. Circadian rhythm of heat production, heart rate, and skin and core temperature under unmasking conditions in men. Am J Physiol 267. doi:10.1152/AJPREGU.1994.267.3.R819

Logan RW, McClung CA. 2018. Rhythms of life: circadian disruption and brain disorders across the lifespan. Nature Reviews Neuroscience 2018 20:1 20:49–65. doi:10.1038/s41583-018-0088-y

Loimaala A, Huikuri H V., Kööbi T, Rinne M, Nenonen A, Vuori I. 2003. Exercise training improves baroreflex sensitivity in type 2 diabetes. Diabetes 52:1837–1842. doi:10.2337/DIABETES.52.7.1837

Maier G, Delezie J, Westermark PO, Santos G, Ritz D, Handschin C. 2022. Transcriptomic, proteomic and phosphoproteomic underpinnings of daily exercise performance and zeitgeber activity of training in mouse muscle. J Physiol 600:769– 796. doi:10.1113/JP281535

Mancilla R, Brouwers B, Schrauwen-Hinderling VB, Hesselink MKC, Hoeks J, Schrauwen P. 2021. Exercise training elicits superior metabolic effects when performed in the afternoon compared to morning in metabolically compromised humans. Physiol Rep 8. doi:10.14814/PHY2.14669

Martin A, Carpentier A, Guissard N, Van Hoecke J, Duchateau J. 1999. Effect of time of day on force variation in a human muscle. Muscle Nerve 22:1380–1387. doi:10.1002/(SICI)1097-4598(199910)22:10

McCarthy JJ, Andrews JL, McDearmon EL, Campbell KS, Barber BK, Miller BH, Walker JR, Hogenesch JB, Takahashi JS, Esser KA. 2007. Identification of the circadian transcriptome in adult mouse skeletal muscle. Physiol Genomics 31:86. doi:10.1152/PHYSIOLGENOMICS.00066.2007

McCurdy CE, Cartee GD. 2005. Akt2 is essential for the full effect of calorie restriction on insulin-stimulated glucose uptake in skeletal muscle. Diabetes 54:1349–1356. doi:10.2337/DIABETES.54.5.1349

Millar-Craig MW, Bishop CN, Raftery EB. 1978. Circadian variation of blood-pressure. Lancet 1:795–797. doi:10.1016/S0140-6736(78)92998-7

Mirizio GG, Nunes RSM, Vargas DA, Foster C, Vieira E. 2020. Time-of-Day Effects on Short-Duration Maximal Exercise Performance. Scientific Reports 2020 10:1 10:1–17. doi:10.1038/s41598-020-66342-w

Molcan L. 2023. Time distributed data analysis by Cosinor. Online application. bioRxiv 805960. doi:10.1101/805960

Nicolas A, Gauthier A, Michaut A, Davenne D. 2007. Effect of circadian rhythm of neuromuscular properties on muscle fatigue during concentric and eccentric isokinetic actions. Isokinet Exerc Sci 15:117–129. doi:10.3233/IES-2007-0258

Peek CB, Affinati AH, Ramsey KM, Kuo HY, Yu W, Sena LA, Ilkayeva O, Marcheva B, Kobayashi Y, Omura C, Levine DC, Bacsik DJ, Gius D, Newgard CB, Goetzman E, Chandel NS, Denu JM, Mrksich M, Bass J. 2013. Circadian clock NAD+ cycle drives mitochondrial oxidative metabolism in mice. Science (1979) **342**. doi:10.1126/SCIENCE.1243417/

Ploug T, Galbo H, Richter EA. 1984. Increased muscle glucose uptake during contractions: no need for insulin. Am J Physiol 247. doi:10.1152/AJPENDO.1984.247.6.E726

Qian J, Xiao Q, Walkup MP, Coday M, Erickson ML, Unick J, Jakicic JM, Hu K, Scheer FAJL, Middelbeek RJW. 2023. Association of Timing of Moderate-to-Vigorous Physical Activity With Changes in Glycemic Control Over 4 Years in Adults With Type 2 Diabetes From the Look AHEAD Trial. Diabetes Care 46:1417–1424. doi:10.2337/DC22-2413

Reduction in the Incidence of Type 2 Diabetes with Lifestyle Intervention or Metformin. 2002. . New England Journal of Medicine 346:393–403. doi:10.1056/NEJMOA012512

Reilly T, Atkinson G, Edwards B, Waterhouse J, Farrelly K, Fairhurst E. 2007. Diurnal variation in temperature, mental and physical performance, and tasks specifically related to football (soccer). Chronobiol Int 24:507–519. doi:10.1080/07420520701420709

Reilly T, Down A. 1986. Circadian Variation in the Standing Broad Jump. Percept Mot Skills 62:830–830. doi:10.2466/PMS.1986.62.3.830

Richter EA, Hargreaves M. 2013. Exercise, GLUT4, and Skeletal Muscle Glucose Uptake. Physiol Rev 93:993–1017. doi:10.1152/physrev.00038.2012.-Glucose

Riddell MC, Turner L V., Patton SR. 2023. Is There an Optimal Time of Day for Exercise? A Commentary on When to Exercise for People Living With Type 1 or Type 2 Diabetes. Diabetes Spectr 36:146. doi:10.2337/DSI22-0017

Rodahl A, O’Brien M, Firth RGR. 1976. Diurnal variation in performance of competitive swimmers. Journal of Sports Medicine and Physical Fitness 16:72–76.

Romijn JA, Coyle EF, Sidossis LS, Gastaldelli A, Horowitz JF, Endert E, Wolfe RR. 1993. Regulation of endogenous fat and carbohydrate metabolism in relation to exercise intensity and duration. Am J Physiol 265. doi:10.1152/AJPENDO.1993.265.3.E380

Sato S, Basse AL, Schönke M, Chen S, Samad M, Altıntaş A, Laker RC, Dalbram E, Barrès R, Baldi P, Treebak JT, Zierath JR, Sassone-Corsi P. 2019. Time of Exercise Specifies the Impact on Muscle Metabolic Pathways and Systemic Energy Homeostasis. Cell Metab 30:92–110.e4. doi:10.1016/J.CMET.2019.03.013

Savikj M, Gabriel BM, Alm PS, Smith J, Caidahl K, Björnholm M, Fritz T, Krook A, Zierath JR, Wallberg-Henriksson H. 2019. Afternoon exercise is more efficacious than morning exercise at improving blood glucose levels in individuals with type 2 diabetes: a randomised crossover trial. Diabetologia 62:233–237. doi:10.1007/S00125-018-4767-Z

Sedliak M, Finni T, Peltonen J, Hakkinen K. 2008. Effect of time-of-day-specific strength training on maximum strength and EMG activity of the leg extensors in men. J Sports Sci 26:1005–1014. doi:10.1080/02640410801930150

Souissi N, Gauthier A, Sesboüé B, Larue J, Davenne D. 2002. Effects of regular training at the same time of day on diurnal fluctuations in muscular performance. J Sports Sci 20:929–937. doi:10.1080/026404102320761813

Stephan FK. 2002. The “other” circadian system: food as a Zeitgeber. J Biol Rhythms 17:284–292. doi:10.1177/074873040201700402

Svensson K, LaBarge SA, Sathe A, Martins VF, Tahvilian S, Cunliffe JM, Sasik R, Mahata SK, Meyer GA, Philp A, David LL, Ward SR, McCurdy CE, Aslan JE, Schenk S. 2020. p300 and cAMP response element-binding protein-binding protein in skeletal muscle homeostasis, contractile function, and survival. J Cachexia Sarcopenia Muscle 11:464. doi:10.1002/JCSM.12522

Um JH, Pendergast JS, Springer DA, Foretz M, Viollet B, Brown A, Kim MK, Yamazaki S, Chung JH. 2011. AMPK Regulates Circadian Rhythms in a Tissue- and Isoform-Specific Manner. PLoS One 6. doi:10.1371/JOURNAL.PONE.0018450

van Moorsel D, Hansen J, Havekes B, Scheer FAJL, Jörgensen JA, Hoeks J, Schrauwen-Hinderling VB, Duez H, Lefebvre P, Schaper NC, Hesselink MKC, Staels B, Schrauwen P. 2016. Demonstration of a day-night rhythm in human skeletal muscle oxidative capacity. Mol Metab 5:635–645. doi:10.1016/J.MOLMET.2016.06.012

Wolff G, Esser KA. 2012. Scheduled exercise phase shifts the circadian clock in skeletal muscle. Med Sci Sports Exerc 44:1663–1670. doi:10.1249/MSS.0B013E318255CF4C

Xin H, Huang R, Zhou M, Chen J, Zhang J, Zhou T, Ji S, Liu X, Tian H, Lam SM, Bao X, Li L, Tong S, Deng F, Shui G, Zhang Z, Wong CCL, Li MD. 2023. Daytime-restricted feeding enhances running endurance without prior exercise in mice. Nat Metab 5:1236–1251. doi:10.1038/S42255-023-00826-7

