## Supplementary figures and images for "Intrinsic skeletal muscle function and contraction-stimulated glucose uptake do not vary by time-of-day in mice"

### Supplementary Figures 1 and 2

##### Supplementary Figure 1

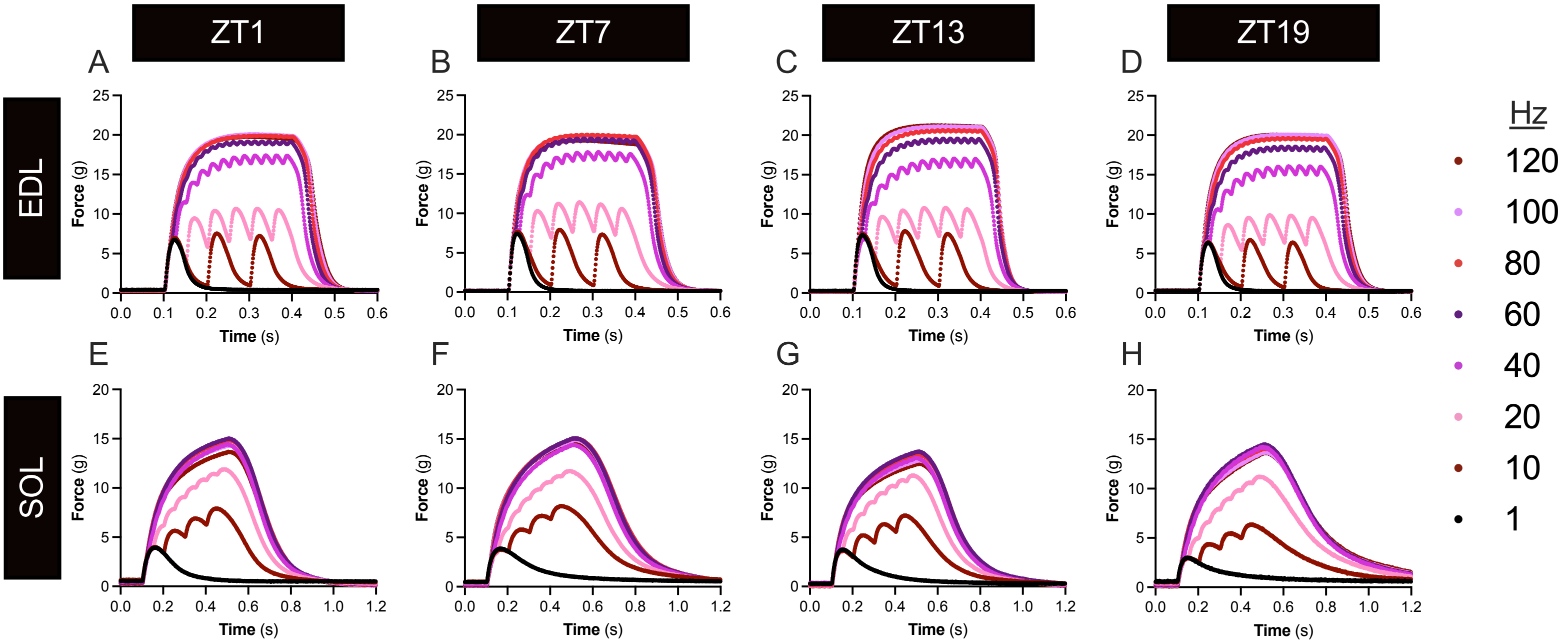

#### Supplementary Figure 2

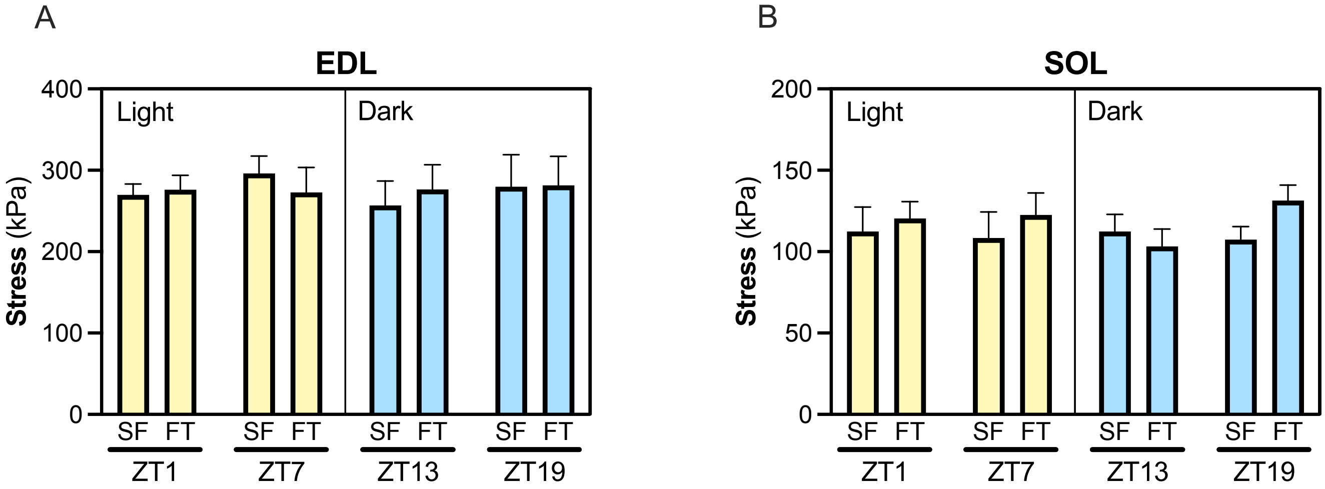
